# Systematic evaluation of integration methods and parameters on single-cell RNA-sequencing biological insights: a case study on cattle embryos

**DOI:** 10.64898/2026.02.01.703145

**Authors:** Fernando Henrique Biase, Michael Morozyuk, Camila Ezepha

## Abstract

**Background:** Single-cell RNA sequencing (scRNA-seq) integration methods remove technical variation while preserving biological signal, yet systematic frameworks for evaluating how parameter choices influence biological interpretation remain limited. Traditional benchmarking approaches evaluate single-parameter configurations per method, potentially missing systematic patterns in functional outcomes and method convergence. A framework for systematic integration parameter evaluation was developed and applied to bovine embryo development.

**Results:** Six integration methods (FastMNN, CCA, RPCA, scVI, Harmony, STACAS) combined with multiple parameters, including those for neighbor identification and clustering, yielded 8232 combinations. The main outputs evaluated were specific cell counts and marker identification. After filtering for extremely poor cell and marker identification, 4,287 integration parameter combinations were retained for analysis. There were three major patterns (clusters) with integration methods distributed non-randomly across clusters and distinct biological outcomes. One pattern emerged, composed of scVI and STACAS integration, dominated by the lack of identification of epiblast cells. Cluster 2 (n=29), also composed of scVI and STACAS integration, identified the most epiblast markers (n=7, 8, or 9) but had a limited number of epiblast cells (median=10). Cluster 1 (n=4,120 combinations) had the highest method diversity. Across clusters, trophoblast and mesoderm showed high functional distinctness, while epiblast and hypoblast showed moderate overlap in gene ontology classes.

**Conclusions:** The approach reveals that parameter choices influence cell type classification, functional interpretation, and the degree of method convergence, with implications for identifying specific biological inferences for further orthogonal validation. A systematic approach to evaluating integration methods, along with other parameters, is advisable for accurate biological inference.

## Background

Single-cell RNA sequencing (scRNA-seq) has transformed developmental biology by enabling comprehensive characterization of cellular heterogeneity during embryogenesis [1–3]. Technical variation between experimental batches necessitates computational integration to enable joint analysis while preserving biological signal [4, 5]. Multiple integration approaches exist, including mutual nearest neighbors (FastMNN) [6], canonical correlation analysis (CCA) [7], reciprocal principal component analysis (RPCA) [8], deep learning (scVI) [9], harmony-based methods [10], and semi-supervised integration (STACAS) [11], each employing distinct algorithmic strategies.

A critical gap in current integration benchmarking is the lack of systematic frameworks for evaluating how parameter choices influence biological interpretation beyond cell type detection. Most benchmarking studies select integration approaches based on computational metrics evaluating batch correction strength [12–14]. Still, these may not capture whether different parameters fundamentally alter biological conclusions regarding lineage-specific functional programs, or whether different integration methods converge on consistent biological interpretations within the same parameter regime. These questions are particularly important in developmental systems where rare but biologically critical populations must be reliably detected, and where functional characterization guides mechanistic hypotheses.

Recent benchmarking studies have compared integration methods [15–17], but most: (1) use default parameters without systematic exploration, (2) evaluate single parameter configurations per method rather than multiple representatives, (3) assess performance using computational rather than biological metrics, and (4) assume methods can be optimized through parameter adjustment without testing whether different methods converge on consistent functional interpretations. The field lacks systematic frameworks for multi-level biological validation of integration parameter choices.

Here, this gap is addressed by developing a framework that enables systematic evaluation of integration parameter effects across multiple biological levels. Our results revealed patterns of cell annotation and marker identification that are highly dependent on the integration method and the cluster identification parameters used. Evaluating one high-performing representative per integration method per pattern enabled the identification of functional divergence from biological results and method convergence. Our framework reveals: (1) parameters result into biological outputs that result into different functional outcomes, (2) specific parameter combinations fail to classify expected cell lineage while other combinations preserve all populations, (3) parameter choices influence both cell classification patterns and functional interpretation, and (4) critically for method selection, different integration methods show varying degrees of functional convergence depending on cell type and biological process, enabling identification of which biological conclusions are method-robust versus requiring additional informatics investigation.

## Methods

An overview of the procedures is depicted on Fig. 1. Overall, single cell RNA-sequencing data was produced from cattle embryo cultured in vitro for either 12 or 16 days (Fig. 1A). Next the data was filtered and processed followed by the development of framework for a systematic evaluation of multiple integration and parameters, including for clustering (Fig. 1B). The output of those analyses were investigated for the formation of patterns of biological signal (Fig. 1C) which were further interrogated for the underlying in silico functional insights (Fig. 1D).

**Fig 1.**
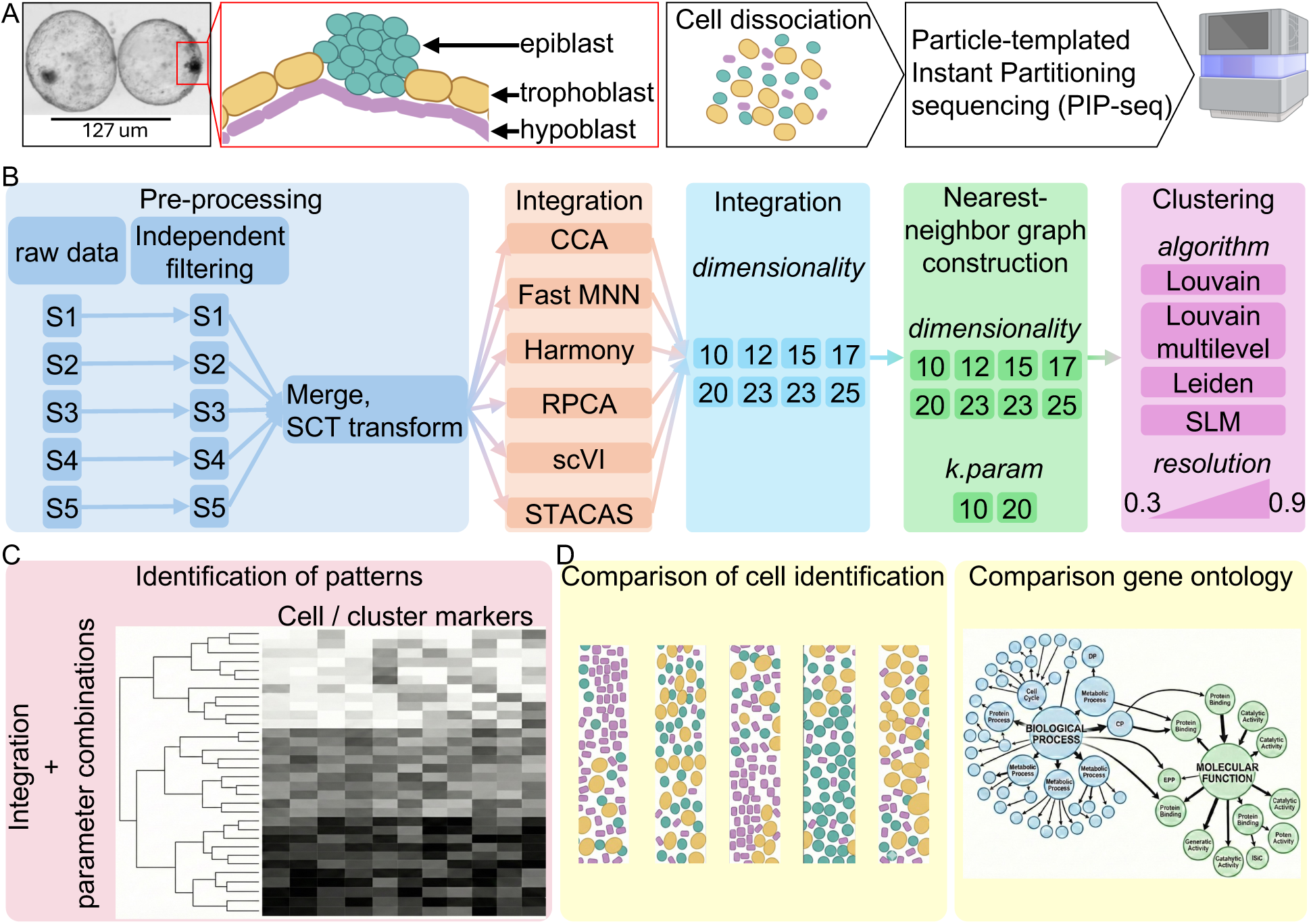
Overview of the procedures. (A) Representative images of the embryos used for the analyses, along with the major expected cell types and the method of PIP-seq used for sequencing. (B) Schematics of the grid search used for the systematic analysis of the data. (C) Schematics of the identification of patterns of the results using hierarchical clustering. (D) Biological insights were obtained from multiple analyses, including cell annotation and in silico functional analysis.

### In vitro embryo production

In vitro embryo production and DNA editing were conducted according to protocols, including media composition, detailed elsewhere [18–20]. Cumulus-oocyte complexes (COCs) were obtained from ovaries sourced from a slaughterhouse. The COCs were selected based on morphology and cultured in oocyte maturation medium in groups of 10 in 50 µL of medium covered by light mineral oil (Sigma-Aldrich, St. Louis, MO). In vitro maturation plates were incubated for 22-24 hours at 38.5 °C and 5 % CO_2_ in a humidified atmosphere. Next, the mature COCs were washed with synthetic oviductal fluid (SOF) medium containing N-2-hydroxyethylpiperazine-N′-2-ethanesulfonic acid (HEPES; Thermo Fisher Scientific) and SOF for fertilization before transferring into a final fertilization plate (100 COCs/mL). Sperm from one bull was prepared from thawed frozen semen straws and processed for in vitro fertilization at a concentration of 1,000,000 spermatozoa/mL. COCs and spermatozoa were co-incubated for approximately 15 hours under the same conditions described for in vitro maturation.

Cumulus cells were removed from the putative zygotes by pulse vortexing in a hyaluronidase (Sigma-Aldrich, St. Louis, MO, 1,400 IU/mL) solution followed by repeated pipetting. Putative zygotes were placed in culture media (25 in 50 µL of synthetic oviduct fluid), covered by mineral oil, and incubated at 38.5 °C with 5 % CO_2_, 5 % O_2_, and 90 % N_2_ in a humidified Eve Benchtop Incubator (WTA, College Station, TX, USA) for approximately 150 hours.

Approximately 150 hours post culture (∼167 hours post fertilization), blastocysts were moved to a post-hatching culture using the media defined elsewhere [21, 22]. One-half of the media was replaced every other day. Embryos were collected on the equivalent days 12 and 16 of *in vitro* culture.

### Single-cell RNA sequencing

Once removed from the media embryos were washed three times in phosphate buffered solution (PBS, Gibco, Grand Island, NY) containing 0.04% of bovine serum albumin (Sigma-Aldrich, St. Louis, MO), and individually placed in 100 µL trypsin solution (TrypLE™ Express Enzyme (1X), no phenol red, Gibco, Grand Island, NY), followed by repeated pipetting with The Stripper™ Pipettor with 150 or 75 µm pipettes (CooperSurgical, Trumbull, CT) for approximately 20 minutes. The trypsin was inactivated by the addition of 100 µL of PBS containing 3% fetal bovine serum (Cytiva, Logan, UT). The cells were then filtered through a 40µm mesh filter (Greiner Bio-One, Kremsmünster, Austria) in 50 mL tube at 200xg for 1 minute. Two hundred mL of solution, including the cell pellet, was eluted in cell buffer from the Illumina Single Cell 3’ RNA Prep, T2 (Illumina, San Diego, CA). Libraries were then prepared as described in the manufacturer’s protocol. This protocol uses particle-templated instant partitioning (PIP-seq) to produce single-cell RNA-sequencing [23]. The sequencing was carried out at The Genomics Sequencing Facility at Virginia Tech using the Illumina NextSeq 2000 equipment.

### Sequence alignment and annotation

The fastq files were processed using the PIPseker^TM^ software for Single Cell Data Analysis, with alignment to the cattle genome (Bos_taurus.ARS-UCD2.0), and fragments counted based on the Bos_taurus.ARS-UCD2.0.114 annotation [24–26].

### Data filtering and processing

The data were imported into RStudio running R version 4.5.2 [27] using Seurat 5.4.0 [7, 8, 28]. Each dataset was filtered based on more than 200 and fewer than 7000 features, which resulted in the retention of data from three embryos collected on day 12 and two embryos collected on day 16. The dataset was merged and transformed using the “SCTransform” function in Seurat. Transformation was carried out using the percentage of mitochondrial genes as a variable for regression. This object was saved and used for all downstream analyses.

### Framework for integration and parameter evaluation

Parameters were systematically varied across six integration methods to generate an initial grid of 8,232 combinations. The methods evaluated were Fast mutual nearest neighbors (FastMNN) [6], canonical correlation analysis (CCA) [7], reciprocal principal component analysis (RPCA) [8], single-cell variational inference (scVI) [9], Harmony: Iterative clustering and correction [10], and semi-supervised anchoring-based integration (STACAS) [11]. For the integrations CCA, FastMNN, RPCA, scVI, and STACAS, we varied the PCA dimensions: 10, 12, 15, 17, 20, 23, and 25. Harmony integration does not use the PCA dimension as a parameter. Next, the shared nearest neighbor graph was calculated using the function “FindNeighbors” in Seurat using clustering dimensions: 10, 12, 15, 17, 20, 23, and 25; and k.param: 10 or 20. The identification of clusters was carried out using the function “FindClusters” in Seurat using the possible algorithms: Leiden, Louvain, Louvain Multilevel, or SLM, and the following resolutions: 0.3, 0.4, 0.5, 0.6, 0.7, 0.8, 0.9. Finally, the function “PrepSCTFindMarkers” was used to prepare the object for testing differential transcript abundance.

### Automated cell annotation

Each of the identified clusters was annotated automatically using the computational platform ScType [29]. The list of genes used for reference was obtained and carefully curated from the literature [30–37]. Cells were classified as either epiblast, hypoblast, trophoblast, mesoderm, or unknown.

### Differential gene transcript abundance

Once the cells were annotated, the Seurat function “FindMarkers” (Wilcoxon Rank Sum test) was used for the identification of markers associated with each cell type (epiblast, hypoblast, trophoblast, or mesoderm) by comparing the transcript abundance of a given cell type versus the remainder of the cells. Using the Seurat functions “AddMetaData” and “AggregateExpression”, a pseudo bulk was created accounting for cell type and each sample analyzed, then used for differential transcription analysis [38]. Similar contrasts as explained above (i.e., epiblast cells versus hypoblast, trophoblast, and mesoderm cells) were created, and two tests were carried out using the pseudo bulk: 1-edgeR Generalized Linear Model Likelihood Ratio Test (glmLRT) [39, 40], and 2-DESeq2 Likelihood ratio test (LRT) [41]. After each test was executed, genes were filtered to retain those with significantly greater transcript abundance in the test cell type (Log_2_(fold change)>0, and false discovery rate [42] adjusted P value < 0.05). Lastly, a list of genes overlapping across all three tests was created for gene ontology tests.

Two outputs were generated from those analyses for each combination of integration and parameters. First, a list of gene markers for each cell was annotated, based on the overlapping results of differential gene transcript tests; and 2-a list of canonical marker genes that overlapped with the marker genes identified in the differential gene transcript tests.

### Identification of patterns through hierarchical clustering

A systematic search for a robust pattern identification of biological outputs was conducted through hierarchical clustering. First, a matrix containing 8232 combinations and related biological information was created containing the following data (Number of Clusters, Epiblast Cells (N), Epiblast Specific Markers, Epiblast Total Markers, Hypoblast Cells (N), Hypoblast Specific Markers, Hypoblast Total Markers, Trophectoderm Cells (N), Trophectoderm Specific Markers, Trophectoderm Total Markers, Mesoderm Cells (N), Mesoderm Specific Markers, Mesoderm Total Markers, Unknown Cells (N), Total Specific Score, Total Overall Markers) (Additional file 1). Next, a filtering was employed to remove extremes using the following criteria to retain data: combinations with ≤75th percentile unknown cells, ≥25th percentile specific score, ≥25th percentile total markers, >2 clusters, and <16 clusters. The biological output mentioned above was used for clustering procedures.

Next, a grid search was conducted to produce multiple dendrograms. The grid used the following distance metrics [43, 44]: canberra, euclidean, manhattan, and maximum, in combination with the following clustering methods [45–48]: average, complete, and ward.D2. Furthermore, the dendrograms were evaluated at multiple numbers of defined clusters (k: 3, 4, 5, 6, 7, 8).

The robustness of each dendrogram was evaluated based on multiple indices. The function “clusterboot” from the package “fpc” was used for nonparametric bootstrap resampling to evaluate cluster-wise stability [49, 50]. The function “NbClust” from the package “NbClust” [51] was used to calculate multiple indices. A final consensus rank was used to select the most stable and separated clustering scheme.

### Statistical analysis of the biological outputs

The independence of distribution of integration methods, as well as the independence of cell type composition across clusters identified, was tested using a chi-square test of independence (𝜒^2^) [52] using the function “chisq.test” from the package “stats”. Cramer’s V was calculated to measure the effect size for the chi-square test of independence using the formula: *V* = 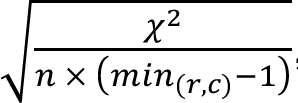, where n= total sample size, min = the minimum value of either the number of rows (r) or columns (c) in the contingency table [53–56]. The proportion of cell composition, as well as the number of canonical markers identified with significantly greater transcript abundance relative to the other cell types, were assessed for differences by the Kruskal-Wallis test [57, 58] using the “kruskal.test” function from the package “stats”. Following a significant Kruskal-Wallis test, pair-wise comparisons were tested with Dunn’s test of multiple comparisons [59] according to the “dunnTest()” function in the package “FSA”. For chi-square, Kruskal-Wallis, and Dunn’s multiple comparisons, significance was inferred at α=0.05.

One combination for each of the integration methods present in each of the three clusters was selected for the enrichment test of gene ontology terms. The list of gene markers for each cell annotated was used for testing. The background used included all genes expressed in the dataset. Gene ontology annotation [60] was obtained from Ensembl [26, 61, 62]. The “goseq” function [63] was used to test enrichment of terms based on the Wallenius non-central hypergeometric distribution [64]. Biological Processes and Molecular Functions were tested separately. False discovery rate [42] was used to adjust the P value, and significance was inferred at FDR <0.05. The similarity between lists of gene ontology terms was compared using the Jaccard index [65].

## Results

### Overview of the dataset

The dataset consisted of single-cell transcriptome of five cattle (*Bos taurus*) embryos produced *in vitro* and collected on days 12 (n=3) and 16 (n=2). Following the sequencing, the data were filtered to retain 1,092 cells with the evaluation of the total of 7,877 genes.

### Distinct cell annotation obtained from the same dataset

Each combination of integration and parameters (Fig. 1B) resulted in the collection of the following data: specific markers for each of the possible cells (epiblast, hypoblast, trophoblast, and mesoderm), total markers identified within the epiblast, hypoblast, trophoblast and mesoderm clusters, along with number of each cell type identified (see Additional file 1 for unfiltered file). The initial question was whether different integration methods and multiple parameters would yield different cell annotations and marker identifications. There was high variability in the number of specific markers identified and in the number of each cell annotated (Additional file 2). Fig. 2 shows the data for specific markers and the number of cells identified. Overall, the number of epiblast markers identified ranged from zero to 11, hypoblast markers ranged from zero to 10, trophectoderm markers ranged from zero to 33, and mesoderm markers ranged from zero to 4 (Fig. 2A, Additional file 2). Harmony integration had the fewest combinations with all cells annotated (0.8%), whereas STACAS had the most (37.2%). The number of cells not annotated ranged from zero to 753 (Fig. 2B). Systematic testing of integration, along with multiple parameters, including clustering, has a critical impact on cell annotation and marker identification.

**Fig. 2.**
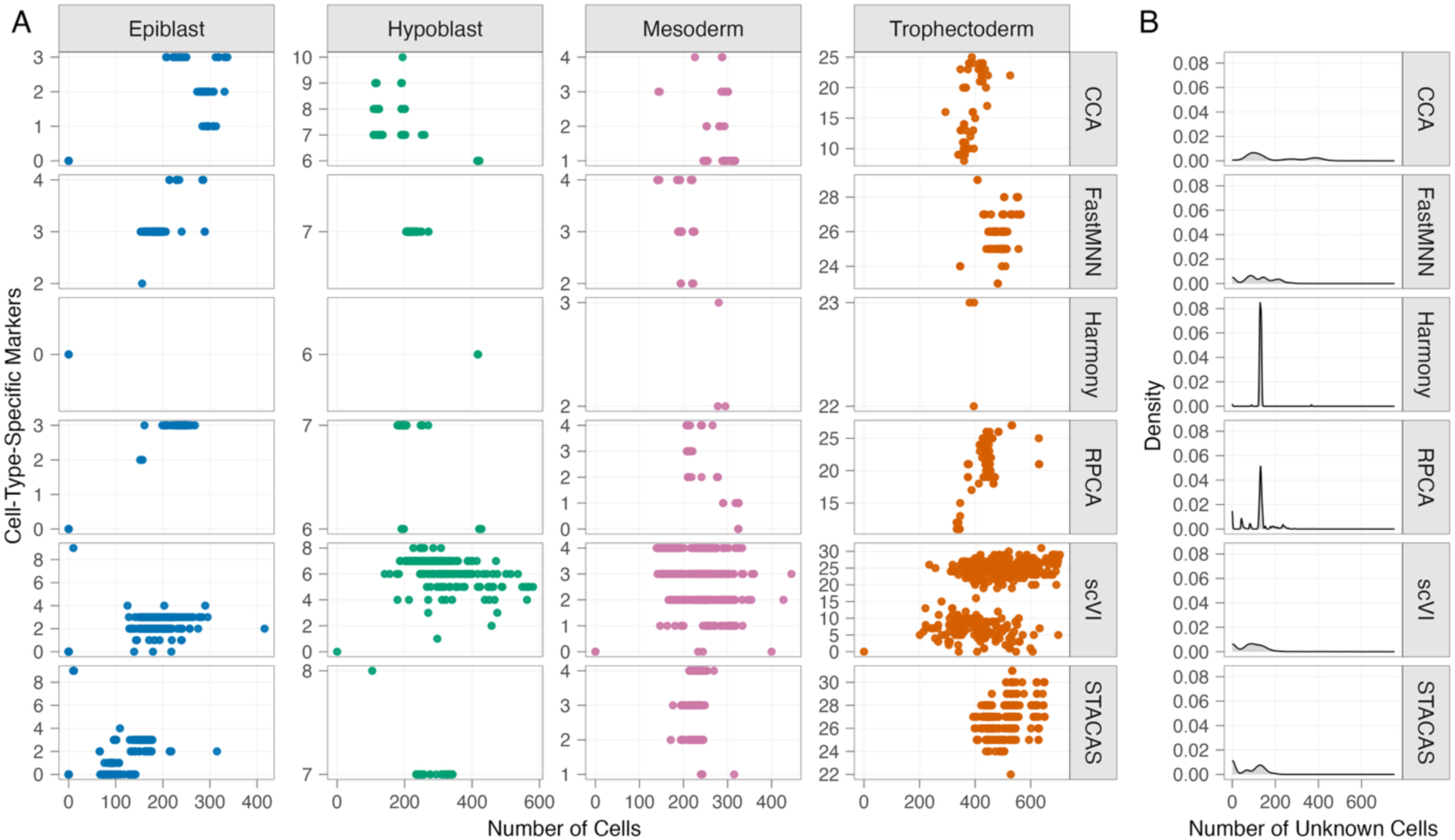
Distribution of annotation and marker identification of multiple integrations and parameters tested. (A) Cell type-specific marker and the number of cells identified in multiple integration methods. Only outcomes that resulted in all cells being annotated were plotted. (B) Distribution of the number of cells not annotated. Additional file 1 presents the results used for graph preparation.

### Distinct patterns of cell annotation resulting from different integration methods

The data with all combinations (Additional file 1) were filtered to remove values that were extreme for specific variables (details in the methods section), resulting in 4,287 combinations used for a systematic search for patterns. Following an unsupervised, systematic search for patterns using hierarchical clustering (details in the methods section), the optimal parameter space for cluster partitioning was achieved using the Canberra distance [66], along with the average linkage [67]. The separation of three clusters resulted in the best combination of indices to indicate a robust patterning identification (Additional file 3). It was possible to determine three distinct and robust patterns in a hierarchical clustering. The three patterns will be referred to as clusters hereafter (Fig 3, see clustering parameters in Additional file 3 and combination assignment in Additional file 4).

**Fig. 3.**
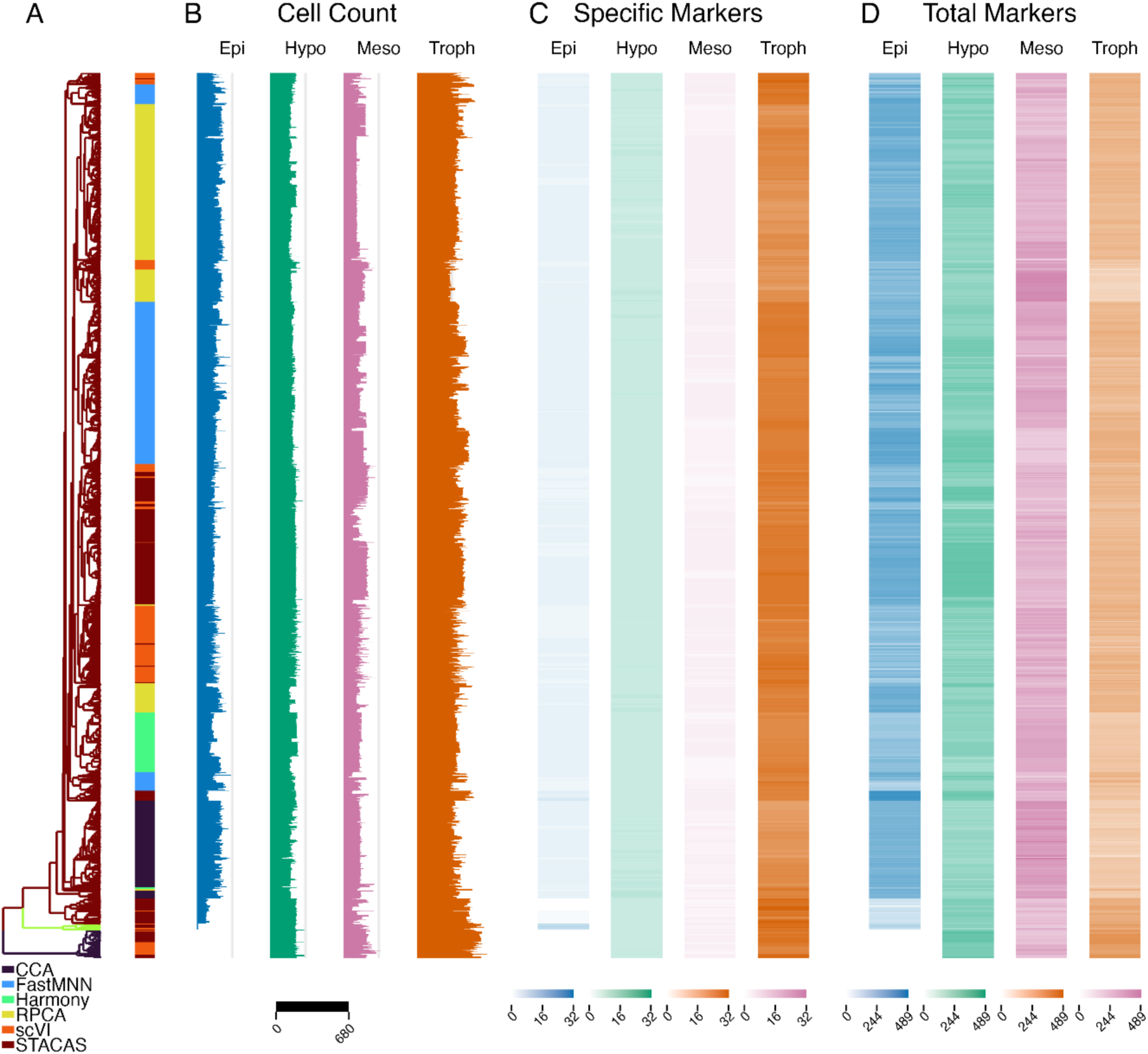
Patterns of biological outcome of multiple combinations of integration and associated parameters. (A) Dendrogram showing the clusters, which are color-coded for k=3. Also present on the panel is the color coding of the six integration methods used. (B) Counts of the four cells annotated on our samples. (C) Number of canonical cell-specific markers identified per cell type. (D) Number of statistically determined markers after the identification of clusters of specific cells.

There was a non-random distribution of integration methods across the clusters (ξ^2^=328.22, P<0.001, Cramér’s V=0.196, Additional file 5). The integration methods CCA, FastMNN, RPCA, and Harmony were only present in Cluster 1. Clusters 2 and 3 contained exclusively STACAS and scVI integration methods. While the method STACAS is highly present in Clusters 2-3, scVI is distributed across all clusters. This suggests method-specific parameter constraints.

### The impact of different integration methods and parameters on cell annotation

The identification of cell types using reference cell markers was one output used for the assessment of the integration, along with multiple parameters tested. Cell type composition differed significantly across clusters (χ^2^=419.77, p<0.001, Cramér’s V=0.259, Fig. 3B, Fig. 4A, Additional file 6). The proportion of all cells was different across all three clusters, with a prominent discrepancy in the identification of epiblast cells. Combinations on Cluster 3 showed no epiblast cells, whereas the combinations in Cluster 2 identified a median of 10 cells, and Cluster 1 identified a median of 196 epiblast cells (H=481.5, P<0.001, Additional file 6). There was also a considerable discrepancy in the identification of trophoblast cells, where Cluster 1 and Cluster 3 annotated a median of 402 and 602, respectively (H=446.07, P<0.001, Additional file 6). Although significant, the number of hypoblast and mesoderm cells annotated showed a less pronounced discrepancy than that of epiblast and trophoblast cells. These results show that integration and the associated tested parameters have a severe impact on cell annotation using canonical markers, thereby influencing cell identification.

**Fig. 4.**
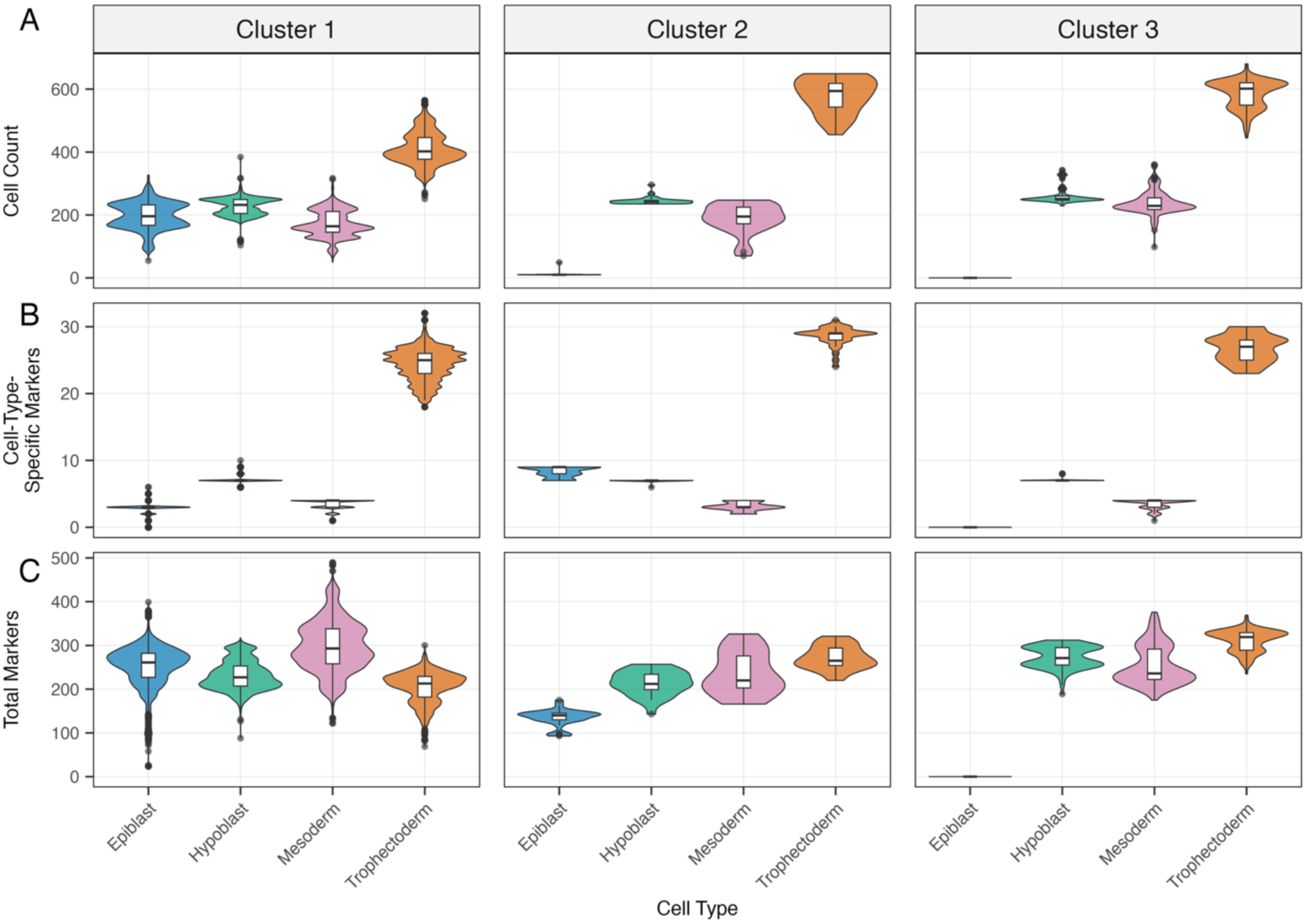
Cell Type Classification and Marker Identification Across Parameter Clusters. Distribution of cell counts (A), cell-type-specific markers (B), and total markers (C) across parameter clusters identified through hierarchical clustering of 4,287 integration parameter combinations.

The second output evaluated was the number of reference markers within the respective cluster of annotated cells supported by differential transcript abundance tests. The number of canonical markers within each cell type was also impacted by the integration method and specific parameters (Fig. 3 C,D, Fig. 4 B,C, Additional file 7). More epiblast canonical markers were identified in Cluster 2 (median 9) than in Clusters 1 and 3 (median 3 and 0, respectively; P<0.001; Fig. 4B, Additional file 7, Additional file 8). A greater number of markers were identified on a small number of epiblast cells (median = 10). Notably, one specific group of 58 combinations (57 STACAS and 1 scVI integration) in cluster 1 had epiblast cells annotated with no canonical markers, with significantly higher transcript abundance than the other cell types. The number of trophoblast canonical markers in Cluster 2 (median 29) was significantly higher than those observed in Clusters 1 and 3 (median 25 and 7, respectively, P<0.001, Fig. 4B, Additional file 7, Additional file 8). The results show that different integration and parameters have an impact on the determination of specific markers.

### The impact of different integration methods and parameters on biological insights

One representative of each integration method within the cluster was selected based on the number of canonical markers within that method (see selected combinations in Additional File 9). The differentially abundant gene transcripts identified by the combination for each cell type were used as test genes for gene ontology analyses. A summary of all significant results of all tests executed is presented in Additional Files 10 and 11. The similarity of gene ontology across multiple integration combinations was assessed using the Jaccard index, which measures the percentage of ontology terms that are shared between two tests relative to the total number of terms annotated.

The degree to which the significant biological process terms overlap across tests was highly variable across cell types. There was no obvious pattern of concordance within clusters or between clusters. The highest concordance achieved for biological processes was observed in epiblast between CCA and RPCA integration in cluster 1 (83% overlapping terms (Fig 5., Fig. 6)). In hypoblast between RPCA in cluster 1 and STACAS in cluster 2 (80% overlapping terms (Fig 5, Fig. 7)). Overall, the average similarity of biological process term was low in trophoblast (mean 27% Fig. 5 and Fig.8), and the lowest in mesoderm (mean 9%, Fig. 5 and Fig. 9). The highest concordance obtained in Molecular function terms was observed in epiblast between Harmony in cluster 1 and scVI in cluster 3 (100% overlapping terms, Fig. 5, Fig 6.), and in trophoblast between CCA and RPCA in cluster 1 (100% overlapping terms, Fig. 5, Fig 8). Trophoblast, nonetheless, showed the lowest average of concordance of molecular function terms across multiple tests (mean 17%, Fig 5, Fig 8). The results show that the choice of integration, along with other parameters, has a significant effect on the biological inferences resultant from in silico functional analysis.

**Fig. 5.**
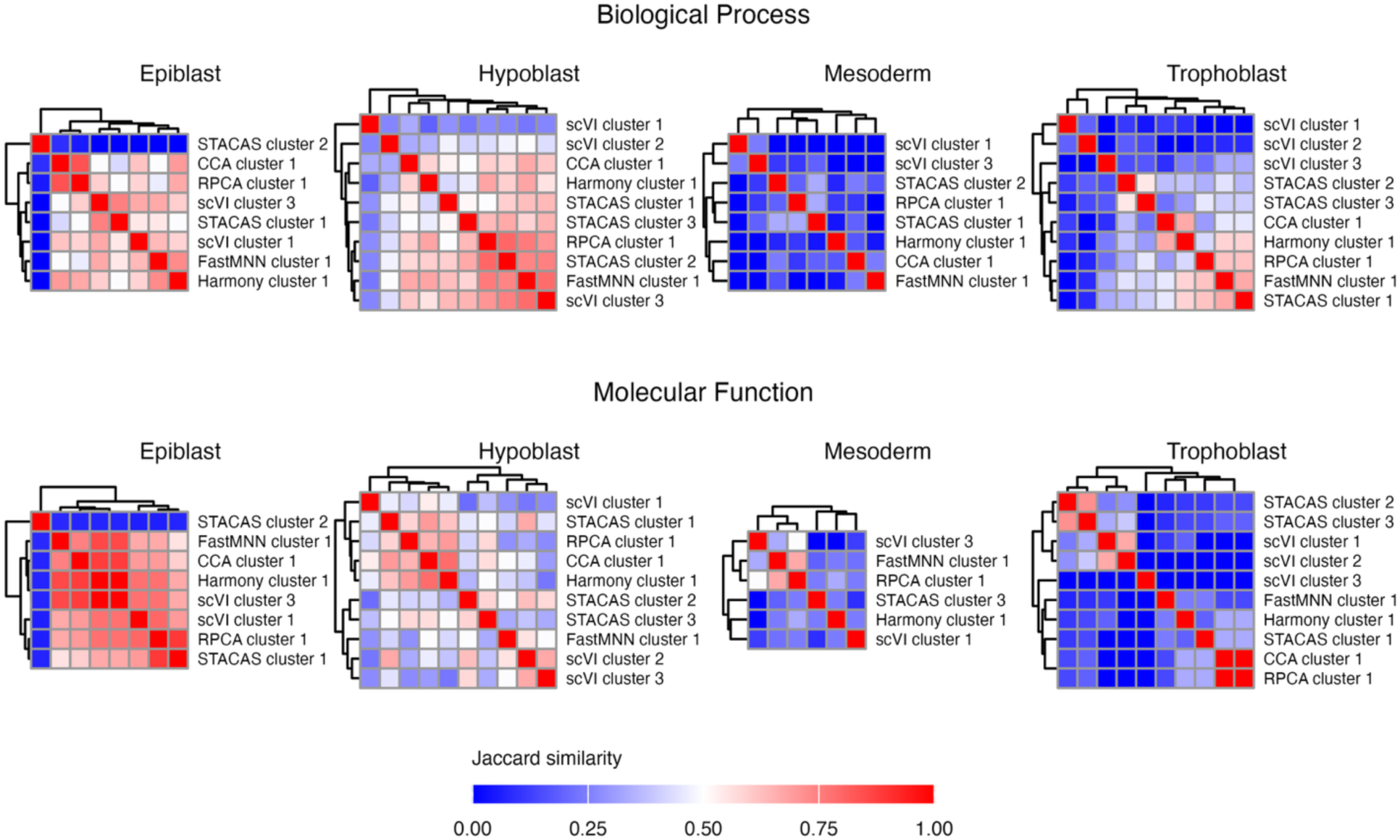
Similarity index (Jaccard) calculated across significantly enriched gene ontology terms resulting from analyses with different integration and parameters.

**Fig 6.**
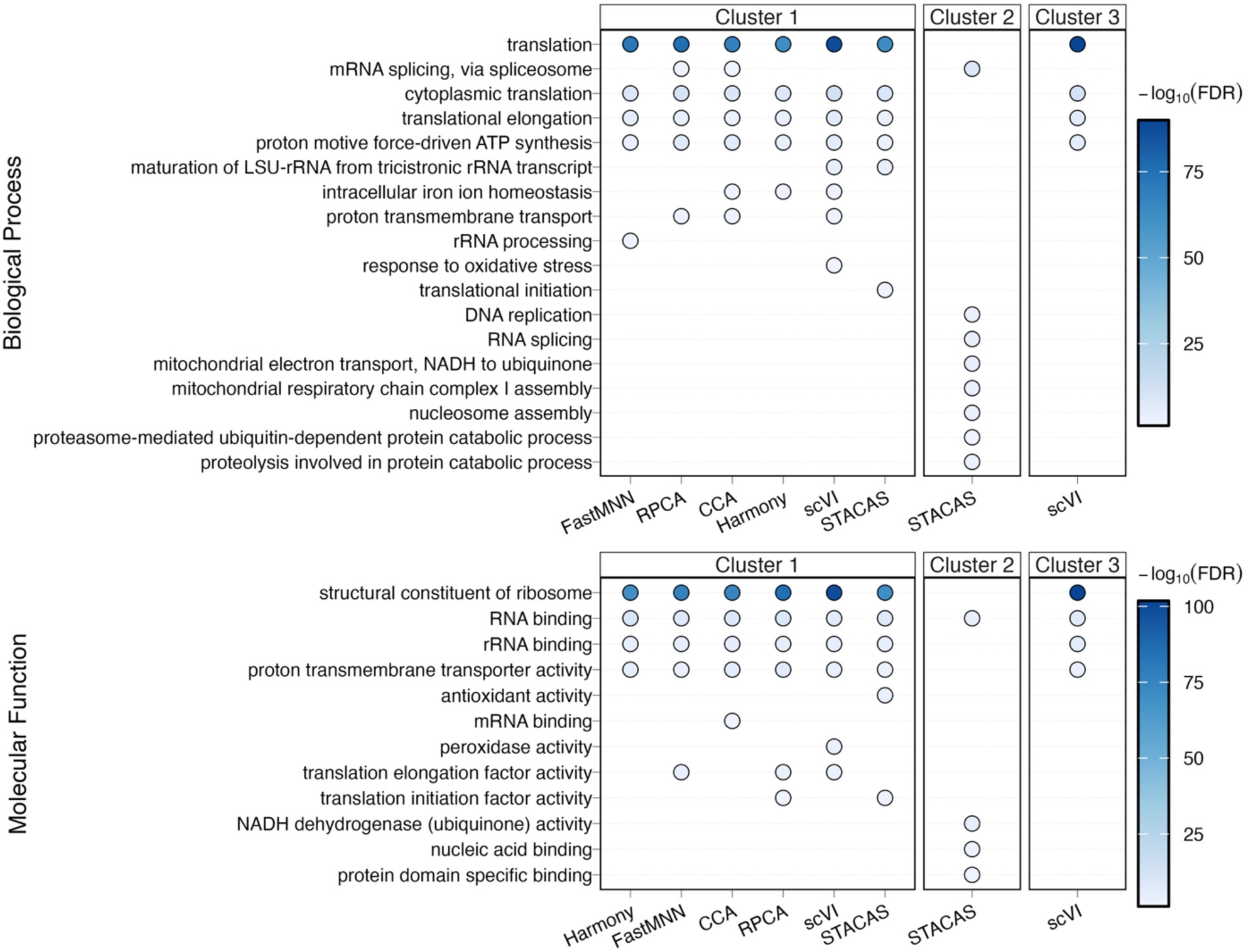
Gene ontology terms enriched in epiblast cells of days 12 and 16 cattle embryos. Only terms with >4 marker genes are displayed for readability. See Additional Files 10 and 11 for the complete list of terms and additional information.

**Fig 7.**
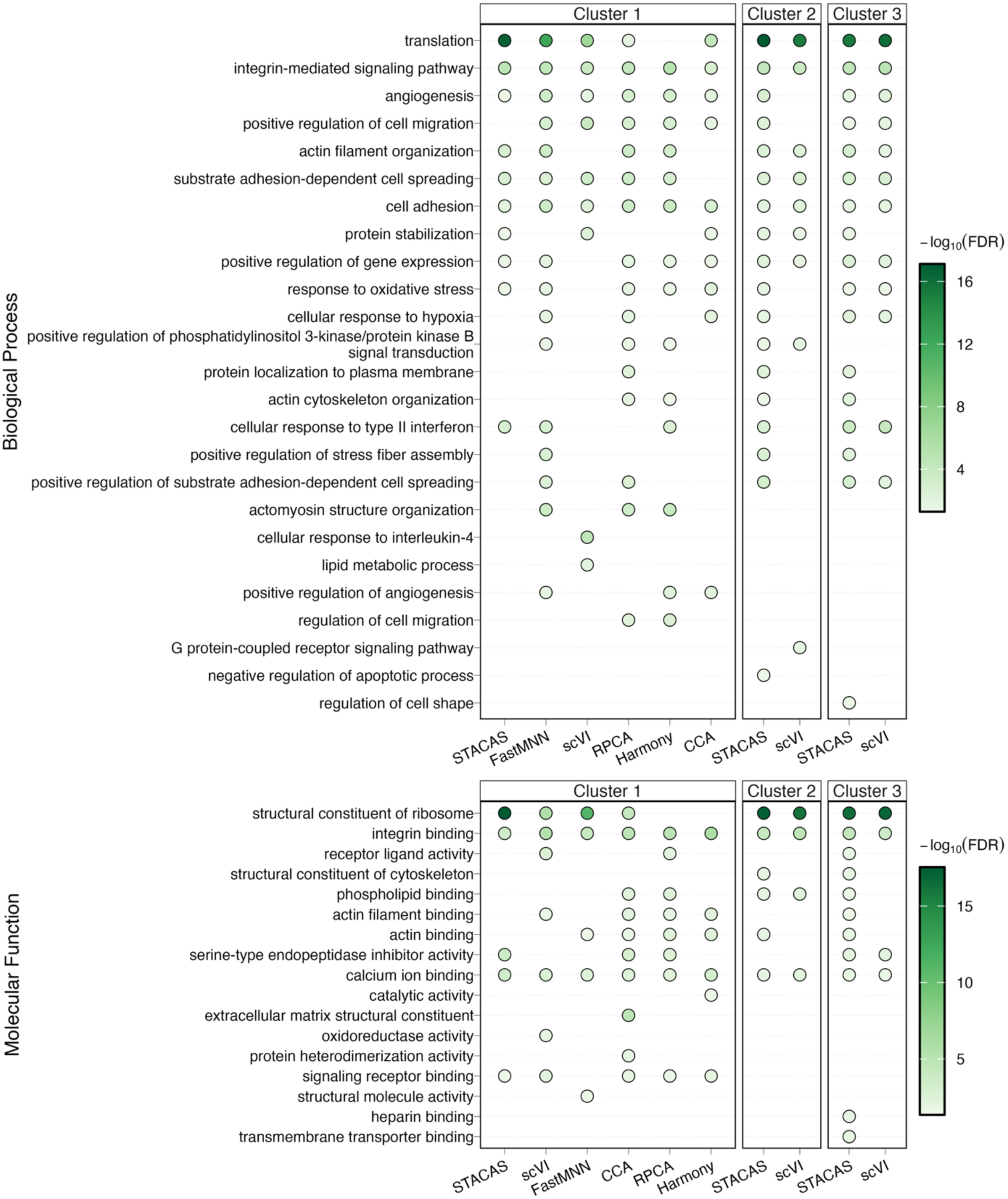
Gene ontology terms enriched in hypoblast cells of days 12 and 16 cattle embryos. Only terms with >4 marker genes are displayed for readability. See Additional Files 10 and 11 for the complete list of terms and additional information.

**Fig 8.**
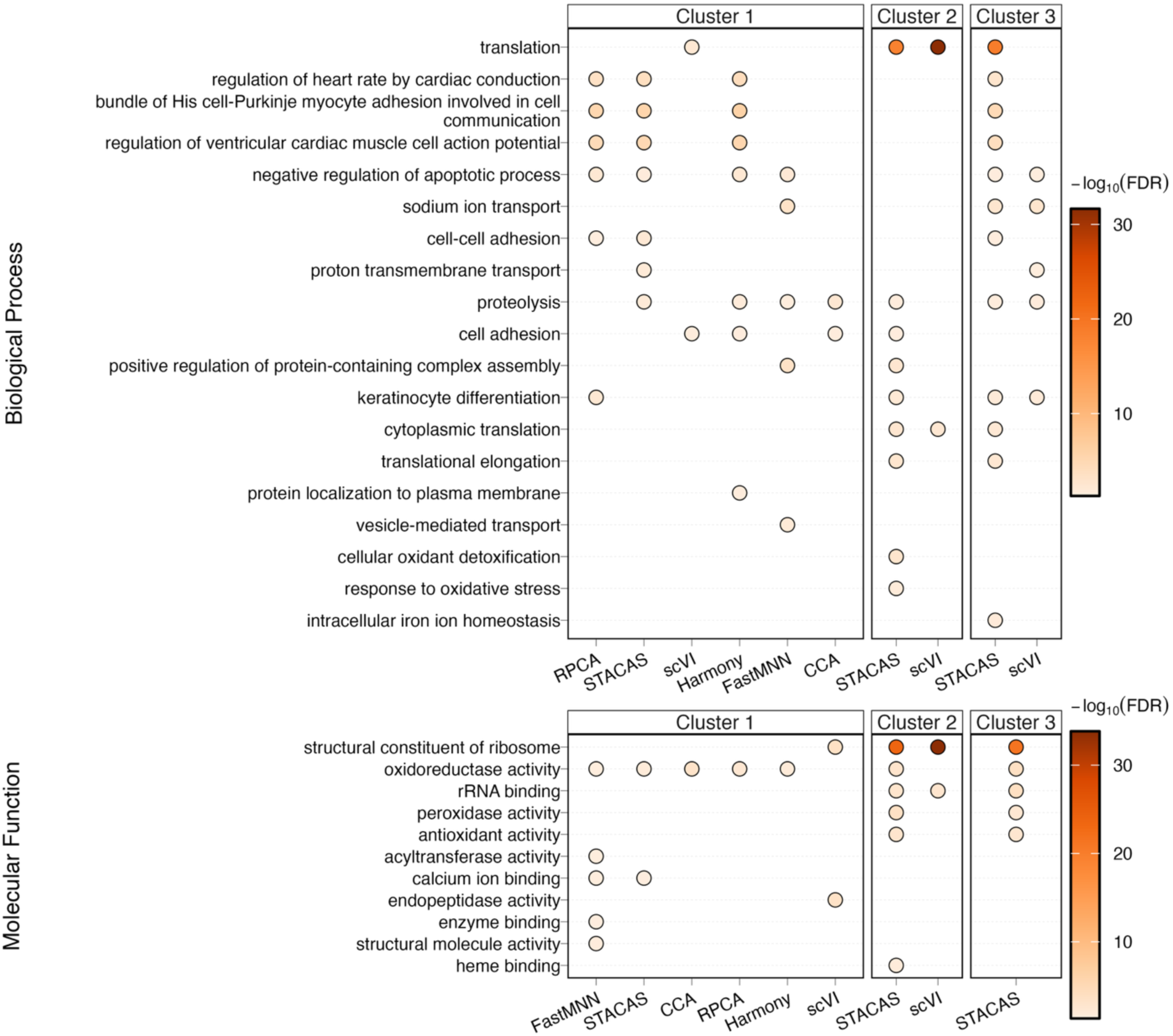
Gene ontology terms enriched in trophoblast cells of days 12 and 16 cattle embryos. Only terms with >4 marker genes are displayed for readability. See Additional Files 10 and 11 for the complete list of terms and additional information.

**Fig 9.**
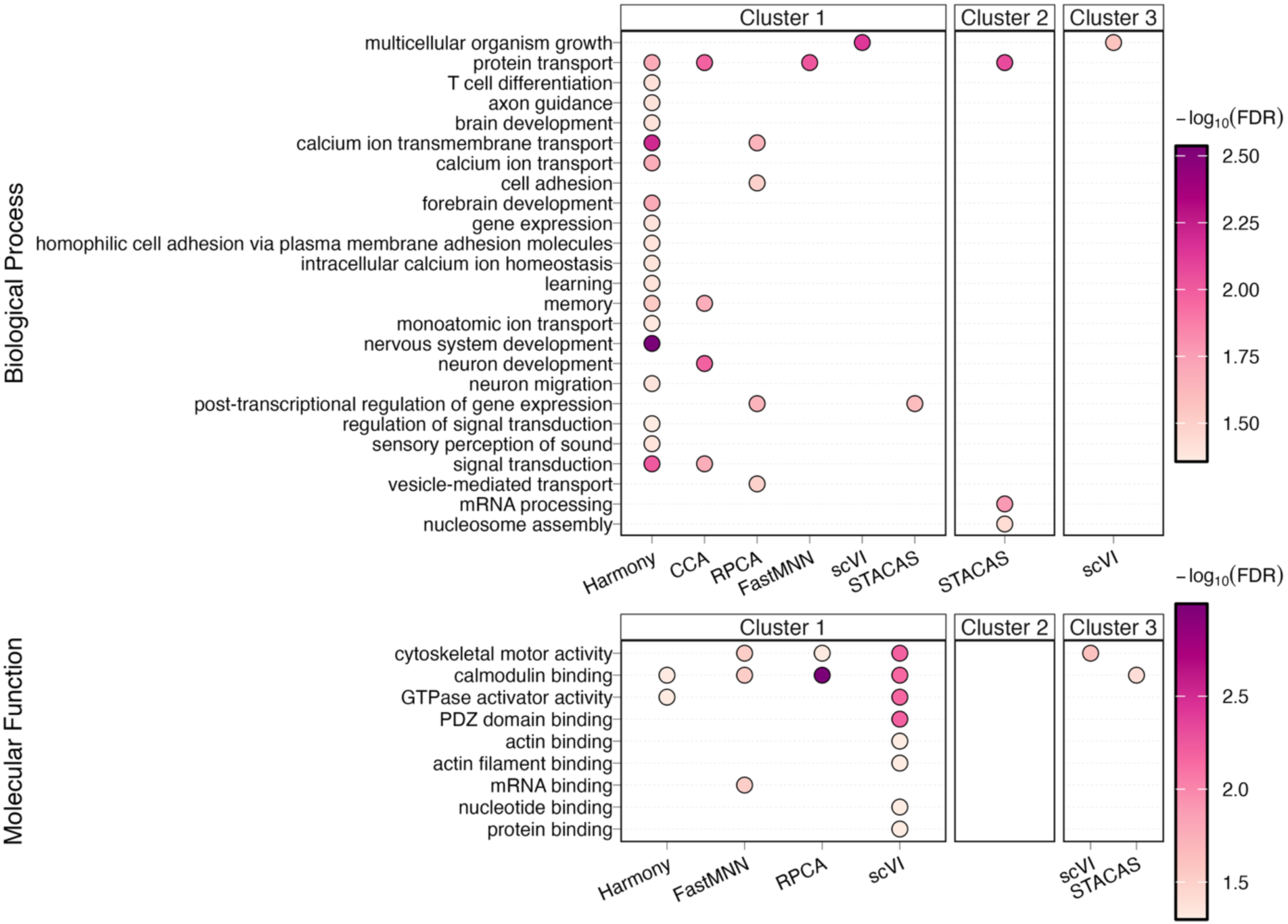
Gene ontology terms enriched in mesoderm cells of day 16 cattle embryos. Only terms with >4 marker genes are displayed for readability. See Additional Files 10 and 11 for the full list of terms and additional information.

## Discussion

The systematic evaluation of integration methods and parameters presented here addresses a critical gap between computational benchmarking and biological validation in single-cell RNA sequencing analysis. Current benchmarking studies predominantly evaluate integration performance using computational metrics such as batch correction strength, graph connectivity, and clustering agreement [5, 15]. Yet these metrics do not directly assess whether analytical choices alter the biological conclusions researchers ultimately draw from their data. The framework developed in this study, testing thousands of parameter combinations and evaluating outputs at multiple biological levels, including cell-type detection, marker identification, and functional enrichment, reveals that the scope of analytical flexibility in scRNA-seq integration is substantially greater than previously appreciated. This finding aligns with growing concerns that parameter choices create “researcher degrees of freedom” [68] capable of yielding contradictory conclusions from identical datasets, as documented across approximately 3,000 pipeline combinations in earlier work evaluating normalization and differential expression choices [69]. The non-random distribution of integration methods across outcome clusters observed here further demonstrates that method-specific constraints interact with parameter settings in ways that cannot be predicted from single-parameter benchmarks alone, consistent with recent findings that no single optimal pipeline exists for all datasets [70].

A particularly consequential finding is that specific method-parameter combinations can prevent detection of biologically critical cell populations. The inability of certain configurations to identify epiblast cells, a lineage essential for understanding embryonic development, illustrates a failure mode that extends beyond the “overcorrection” phenomenon documented in previous integration benchmarks [11, 71]. While prior studies have shown that rare cell populations comprising less than 1% of datasets are difficult for any clustering method to detect reliably [72], the present results demonstrate that even moderately abundant [21, 22] populations can be lost entirely depending on integration and clustering parameter choices. This observation carries significant implications for developmental biology studies, where the biological question often centers on lineage-specification events involving transiently appearing or low-abundance cell types. Researchers relying on default parameters or single integration methods risk drawing conclusions from analyses that have systematically excluded the very populations of greatest biological interest, a concern amplified by evidence that the commonly used Seurat resolution default of 0.8 may not be optimal for most datasets [73]. The finding that some combinations identified more canonical markers within fewer cells, while others identified more cells with fewer supporting markers, further highlights a sensitivity-specificity trade-off that is not captured by standard benchmarking metrics focused on batch mixing and cluster separation.

The evaluation of functional enrichment concordance across integration approaches reveals an underexplored dimension of analytical variability that persists even when cell clustering appears similar. The observation that gene ontology similarity varied dramatically across cell types, with some lineages showing high concordance between methods while others showing minimal overlap, indicates that the choice of integration method and parameters propagates throughout the analytical workflow, affecting biological interpretation. This finding resonates with recent benchmarking of 46 differential expression workflows, which showed that different analytical pipelines detect entirely different pathway categories from the same data, and batch correction can introduce artifacts into fold-change values used for enrichment analysis [74]. Furthermore, functional analysis tool performance is more sensitive to gene set composition than to the statistical method employed [75], suggesting that upstream integration choices that alter marker gene identification will necessarily affect downstream functional conclusions. The cell-type-specific nature of functional concordance observed here suggests that certain biological processes may be more robust to analytical choices than others. However, the factors determining this robustness remain unclear. For studies where functional characterization guides mechanistic hypotheses or identifies therapeutic targets, low concordance across reasonable analytical approaches raises fundamental questions about which biological conclusions warrant confidence and which require orthogonal validation.

The patterns of method convergence and divergence documented in this study have practical implications for how researchers should approach the selection and validation of integration methods. The observation that some integration methods clustered together in outcome space while others were distributed across multiple patterns suggests that certain methods may be more sensitive to parameter perturbations than others, consistent with reports that adjusting clustering parameters remains ad hoc and poorly understood, with poorly understood effects on accuracy and reproducibility [76]. Notably, the methods showing the broadest distribution across outcome patterns may offer greater flexibility but also greater risk of parameter-dependent conclusions. In contrast, methods restricted to specific outcome patterns may be more consistent but potentially biased toward particular types of solutions. These findings support emerging recommendations for ensemble approaches that assess the robustness of conclusions across multiple methods [4, 77], rather than relying on a single “best” method identified from computational benchmarks. The comprehensive scIB benchmark notably found that no method universally outperformed others across all scenarios [15], and the present study extends this observation by demonstrating that method performance also varies across biological output types within a single dataset. The framework presented here provides a template for such multi-method validation by enabling systematic identification of which biological conclusions are method-robust versus parameter-sensitive.

## Conclusions

In conclusion, this study demonstrates that integration method and parameter choices in scRNA-seq analysis affect not only cell clustering but also cell type detection, marker identification, and functional interpretation in ways that current benchmarking approaches do not fully capture. The systematic grid-search framework developed here enables researchers to quantify analytical flexibility and identify conditions under which different methods converge on consistent biological conclusions. Future work should extend this framework to additional biological systems to determine whether the patterns of method-specific behavior observed in embryonic development generalize across tissue types and experimental designs. Additionally, the development of guidelines specifying which parameter combinations are inappropriate for specific analytical objectives, analogous to the filtering criteria applied here, would help researchers avoid configurations that systematically fail to detect expected cell populations. As single-cell technologies become increasingly central to biological discovery, ensuring that analytical choices do not predetermine biological conclusions represents an essential step toward reproducible and reliable inference from these powerful datasets.

## Declarations

### Ethics approval and consent to participate

Not applicable

### Consent for publication

Not applicable

### Availability of data and material

The datasets used and/or analyzed during the current study are available from the corresponding author on reasonable request.

### Authors’ contributions

FB contributed to data acquisition, data analysis, interpretation, and writing of the manuscript; CE and MM contributed to the data acquisition and draft revision. All authors read and approved the final manuscript.

## Acknowledgements

We thank the Genomics Sequencing Facility at Virginia Tech Fralin Life Sciences Institute for the support sequencing the libraries prepared.

## List of Abbreviations

scRNA-seq: Single-cell RNA sequencing
PIP-seq: Particle-templated instant partitioning (sequencing method)
FastMNN: Fast mutual nearest neighbors
CCA: Canonical correlation analysis
RPCA: Reciprocal principal component analysis
scVI: Single-cell variational inference
STACAS: Semi-supervised anchoring-based integration (Sub-Type Anchor Correction for Alignment in Seurat)
PCA: Principal component analysis
COCs: Cumulus-oocyte complexes
SOF: Synthetic oviductal fluid
HEPES: N-2-hydroxyethylpiperazine-N′-2-ethanesulfonic acid
PBS: Phosphate buffered solution
glmLRT: Generalized Linear Model Likelihood Ratio Test
LRT: Likelihood ratio test
FDR: False discovery rate
SLM: Smart Local Moving (clustering algorithm)
GO: Gene ontology

## References

1. Pijuan-Sala B, Griffiths JA, Guibentif C, Hiscock TW, Jawaid W, Calero-Nieto FJ, et al. A single-cell molecular map of mouse gastrulation and early organogenesis. Nature. 2019;566(7745):490–5. 10.1038/s41586-019-0933-9

2. Wagner DE, Klein AM. Lineage tracing meets single-cell omics: opportunities and challenges. Nat Rev Genet. 2020;21(7):410–27. 10.1038/s41576-020-0223-2

3. Tanay A, Regev A. Scaling single-cell genomics from phenomenology to mechanism. Nature. 2017;541(7637):331–8. 10.1038/nature21350

4. Luecken MD, Theis FJ. Current best practices in single-cell RNA-seq analysis: a tutorial. Mol Syst Biol. 2019;15(6):e8746. 10.15252/msb.20188746

5. Tran HTN, Ang KS, Chevrier M, Zhang X, Lee NYS, Goh M, et al. A benchmark of batch-effect correction methods for single-cell RNA sequencing data. Genome Biol. 2020;21(1):12. 10.1186/s13059-019-1850-9

6. Haghverdi L, Lun ATL, Morgan MD, Marioni JC. Batch effects in single-cell RNA-sequencing data are corrected by matching mutual nearest neighbors. Nat Biotechnol. 2018;36(5):421–7. 10.1038/nbt.4091

7. Butler A, Hoffman P, Smibert P, Papalexi E, Satija R. Integrating single-cell transcriptomic data across different conditions, technologies, and species. Nat Biotechnol. 2018;36(5):411–20. 10.1038/nbt.4096

8. Stuart T, Butler A, Hoffman P, Hafemeister C, Papalexi E, Mauck WM, 3rd, et al. Comprehensive Integration of Single-Cell Data. Cell. 2019;177(7):1888–902 e21. 10.1016/j.cell.2019.05.031

9. Lopez R, Regier J, Cole MB, Jordan MI, Yosef N. Deep generative modeling for single-cell transcriptomics. Nat Methods. 2018;15(12):1053–8. 10.1038/s41592-018-0229-2

10. Korsunsky I, Millard N, Fan J, Slowikowski K, Zhang F, Wei K, et al. Fast, sensitive and accurate integration of single-cell data with Harmony. Nat Methods. 2019;16(12):1289–96. 10.1038/s41592-019-0619-0

11. Andreatta M, Carmona SJ. STACAS: Sub-Type Anchor Correction for Alignment in Seurat to integrate single-cell RNA-seq data. Bioinformatics. 2021;37(6):882–4. 10.1093/bioinformatics/btaa755

12. Hou R, Denisenko E, Forrest ARR. scMatch: a single-cell gene expression profile annotation tool using reference datasets. Bioinformatics. 2019;35(22):4688–95. 10.1093/bioinformatics/btz292

13. Mereu E, Lafzi A, Moutinho C, Ziegenhain C, McCarthy DJ, Alvarez-Varela A, et al. Benchmarking single-cell RNA-sequencing protocols for cell atlas projects. Nat Biotechnol. 2020;38(6):747–55. 10.1038/s41587-020-0469-4

14. Chazarra-Gil R, van Dongen S, Kiselev VY, Hemberg M. Flexible comparison of batch correction methods for single-cell RNA-seq using BatchBench. Nucleic Acids Res. 2021;49(7):e42. 10.1093/nar/gkab004

15. Luecken MD, Buttner M, Chaichoompu K, Danese A, Interlandi M, Mueller MF, et al. Benchmarking atlas-level data integration in single-cell genomics. Nat Methods. 2022;19(1):41–50. 10.1038/s41592-021-01336-8

16. Argelaguet R, Cuomo ASE, Stegle O, Marioni JC. Computational principles and challenges in single-cell data integration. Nat Biotechnol. 2021;39(10):1202–15. 10.1038/s41587-021-00895-7

17. Lahnemann D, Koster J, Szczurek E, McCarthy DJ, Hicks SC, Robinson MD, et al. Eleven grand challenges in single-cell data science. Genome Biol. 2020;21(1):31. 10.1186/s13059-020-1926-6

18. Biase FH, Schettini G. Protocol for the electroporation of CRISPR-Cas for DNA and RNA targeting in Bos taurus zygotes. STAR Protoc. 2024;5(1):102940. 10.1016/j.xpro.2024.102940

19. Nix JL, Schettini GP, Speckhart SL, Ealy AD, Biase FH. Ablation of OCT4 function in cattle embryos by double electroporation of CRISPR-Cas for DNA and RNA targeting (CRISPR-DART). PNAS Nexus. 2023;2(11):pgad343. 10.1093/pnasnexus/pgad343

20. Nix J, Marrella MA, Oliver MA, Ealy AD, Biase FH. Cleavage Kinetics is a Better Indicator of Embryonic Developmental Competency Than Brilliant Cresyl Blue Staining of Oocytes. Anim Reprod Sci. 2022:107174.

21. Ramos-Ibeas P, Perez-Gomez A, Gonzalez-Brusi L, Quiroga AC, Bermejo-Alvarez P. Pre-hatching exposure to N2B27 medium improves post-hatching development of bovine embryos in vitro. Theriogenology. 2023;205:73–8. 10.1016/j.theriogenology.2023.04.018

22. Ramos-Ibeas P, Lamas-Toranzo I, Martinez-Moro A, de Frutos C, Quiroga AC, Zurita E, et al. Embryonic disc formation following post-hatching bovine embryo development in vitro. Reproduction. 2020;160(4):579–89. 10.1530/REP-20-0243

23. Clark IC, Fontanez KM, Meltzer RH, Xue Y, Hayford C, May-Zhang A, et al. Microfluidics-free single-cell genomics with templated emulsification. Nat Biotechnol. 2023;41(11):1557–66. 10.1038/s41587-023-01685-z

24. Flicek P, Amode MR, Barrell D, Beal K, Billis K, Brent S, et al. Ensembl 2014. Nucleic Acids Res. 2014;42(Database issue):D749–55. 10.1093/nar/gkt1196

25. Flicek P, Amode MR, Barrell D, Beal K, Brent S, Carvalho-Silva D, et al. Ensembl 2012. Nucleic Acids Res. 2012;40:D84–90. 10.1093/nar/gkr991

26. Kinsella RJ, Kähäri A, Haider S, Zamora J, Proctor G, Spudich G, et al. Ensembl BioMarts: a hub for data retrieval across taxonomic space. Database. 2011;2011:bar030. 10.1093/database/bar030

27. RCoreTeam. R: A Language and Environment for Statistical Computing. Vienna, Austria: R Foundation for Statistical Computing; 2020.

28. Hao Y, Stuart T, Kowalski MH, Choudhary S, Hoffman P, Hartman A, et al. Dictionary learning for integrative, multimodal and scalable single-cell analysis. Nat Biotechnol. 2024;42(2):293–304. 10.1038/s41587-023-01767-y

29. Ianevski A, Giri AK, Aittokallio T. Fully-automated and ultra-fast cell-type identification using specific marker combinations from single-cell transcriptomic data. Nat Commun. 2022;13(1):1246. 10.1038/s41467-022-28803-w

30. Scatolin GN, Ming H, Wang Y, Iyyappan R, Gutierrez-Castillo E, Zhu L, et al. Single-cell transcriptional landscapes of bovine peri-implantation development. iScience. 2024;27(4):109605. 10.1016/j.isci.2024.109605

31. Mole MA, Coorens THH, Shahbazi MN, Weberling A, Weatherbee BAT, Gantner CW, et al. A single cell characterisation of human embryogenesis identifies pluripotency transitions and putative anterior hypoblast centre. Nat Commun. 2021;12(1):3679. 10.1038/s41467-021-23758-w

32. Jia GX, Ma WJ, Wu ZB, Li S, Zhang XQ, He Z, et al. Single-cell transcriptomic characterization of sheep conceptus elongation and implantation. Cell Rep. 2023;42(8):112860. 10.1016/j.celrep.2023.112860

33. Zhao C, Plaza Reyes A, Schell JP, Weltner J, Ortega NM, Zheng Y, et al. A comprehensive human embryo reference tool using single-cell RNA-sequencing data. Nat Methods. 2025;22(1):193–206. 10.1038/s41592-024-02493-2

34. Degrelle SA, Liu F, Laloe D, Richard C, Le Bourhis D, Rossignol MN, et al. Understanding bovine embryo elongation: a transcriptomic study of trophoblastic vesicles. Front Physiol. 2024;15:1331098. 10.3389/fphys.2024.1331098

35. Hue I, Dufort I, Vitorino Carvalho A, Laloe D, Peynot N, Degrelle SA, et al. Different pre-implantation phenotypes of bovine blastocysts produced in vitro. Reproduction. 2018;157(2):163–78. 10.1530/REP-18-0439

36. Hue I, Evain-Brion D, Fournier T, Degrelle SA. Primary Bovine Extra-Embryonic Cultured Cells: A New Resource for the Study of In Vivo Peri-Implanting Phenotypes and Mesoderm Formation. PLoS One. 2015;10(6):e0127330. 10.1371/journal.pone.0127330

37. Hu C, Li T, Xu Y, Zhang X, Li F, Bai J, et al. CellMarker 2.0: an updated database of manually curated cell markers in human/mouse and web tools based on scRNA-seq data. Nucleic Acids Res. 2023;51(D1):D870–D6. 10.1093/nar/gkac947

38. Squair JW, Gautier M, Kathe C, Anderson MA, James ND, Hutson TH, et al. Confronting false discoveries in single-cell differential expression. Nat Commun. 2021;12(1):5692. 10.1038/s41467-021-25960-2

39. Lun ATL, Chen YS, Smyth GK. It’s DE-licious: A recipe for differential expression analyses of RNA-seq experiments using quasi-likelihood methods in edgeR. Statistical Genomics: Methods and Protocols. 2016;1418:391–416. 10.1007/978-1-4939-3578-9_19

40. McCarthy DJ, Smyth GK. edgeR: a Bioconductor package for differential expression analysis of digital gene expression data. Bioinformatics. 2010;26:139–40. 10.1093/bioinformatics/btp616

41. Love MI, Huber W, Anders S. Moderated estimation of fold change and dispersion for RNA-seq data with DESeq2. Genome Biol. 2014;15:550. 10.1186/PREACCEPT-8897612761307401

42. Benjamini Y, Hochberg Y. Controlling the false discovery rate - a practical and powerful approach to multiple testing. J Roy Stat Soc B Met. 1995;57(1):289–300.

43. Mardia KV, Kent JT, Taylor CC. Multivariate analysis: John Wiley & Sons; 2024.

44. Borg I, Groenen PJ. Modern multidimensional scaling: Theory and applications: Springer; 2005.

45. Anderberg MR, editor Cluster Analysis for Applications1973.

46. Hartigan JA. Clustering Algorithms: Wiley; 1975.

47. Murtagh F, Contreras P. Algorithms for hierarchical clustering: an overview, II. Wiley Interdisciplinary Reviews: Data Mining and Knowledge Discovery. 2017;7(6):e1219.

48. Murtagh F, Contreras P. Algorithms for hierarchical clustering: an overview. Wiley interdisciplinary reviews: data mining and knowledge discovery. 2012;2(1):86–97.

49. Hennig C. Dissolution point and isolation robustness: robustness criteria for general cluster analysis methods. Journal of multivariate analysis. 2008;99(6):1154–76.

50. Hennig C. Cluster-wise assessment of cluster stability. Comput Stat Data Anal. 2007;52:258–71.

51. Charrad M, Ghazzali N, Boiteau V, Niknafs A. NbClust: An R Package for Determining the Relevant Number of Clusters in a Data Set. J Stat Softw. 2014;61(6):1–36. 10.18637/jss.v061.i06

52. Pearson K. X. On the criterion that a given system of deviations from the probable in the case of a correlated system of variables is such that it can be reasonably supposed to have arisen from random sampling. The London, Edinburgh, and Dublin Philosophical Magazine and Journal of Science. 1900;50(302):157–75. 10.1080/14786440009463897

53. Cohen J. Statistical Power Analysis for the Behavioral Sciences. Second Edition ed. New York: Taylor & Francis Group; 1988.

54. Yule GU. On the methods of measuring association between two attributes. Journal of the Royal Statistical Society. 1912;75(6):579–652.

55. Pearson K, Heron D. On theories of association. Biometrika. 1913;9(1/2):159–315.

56. Berry KJ, Johnston JE, Mielke Jr PW. A measure of effect size for R× C contingency tables. Psychological reports. 2006;99(1):251–6.

57. Kloke J, McKean J. Nonparametric statistical methods using R: Chapman and Hall/CRC; 2024.

58. Hollander M, Wolfe DA, Chicken E. Nonparametric statistical methods: John Wiley & Sons; 2013.

59. Dunn OJ. Multiple Comparisons Using Rank Sums. Technometrics. 1964;6(3):241–52. 10.2307/1266041

60. Ashburner M, Ball CA, Blake JA, Botstein D, Butler H, Cherry JM, et al. Gene ontology: tool for the unification of biology. The Gene Ontology Consortium. Nat Genet. 2000;25(1):25–9. 10.1038/75556

61. Durinck S, Spellman PT, Birney E, Huber W. Mapping identifiers for the integration of genomic datasets with the R/Bioconductor package biomaRt. Nat Protoc. 2009;4(8):1184–91. 10.1038/nprot.2009.97

62. Durinck S, Moreau Y, Kasprzyk A, Davis S, De Moor B, Brazma A, et al. BioMart and Bioconductor: a powerful link between biological databases and microarray data analysis. Bioinformatics. 2005;21(16):3439–40. 10.1093/bioinformatics/bti525

63. Young MD, Wakefield MJ, Smyth GK, Oshlack A. Gene ontology analysis for RNA-seq: accounting for selection bias. Genome Biol. 2010;11(2):R14. 10.1186/gb-2010-11-2-r14

64. Fog A. Calculation Methods for Wallenius’ Noncentral Hypergeometric Distribution. Communications in Statistics - Simulation and Computation. 2008;37(2):258–73. 10.1080/03610910701790269

65. Gan M. Correlating information contents of gene ontology terms to infer semantic similarity of gene products. Comput Math Methods Med. 2014;2014:891842. 10.1155/2014/891842

66. Lance GN, Williams WT. A general theory of classificatory sorting strategies: II. Clustering systems. The Computer Journal. 1967;10(3):271–7. 10.1093/comjnl/10.3.271

67. Murtagh F. Multidimensional clustering algorithms: Physica; 1985.

68. Simmons JP, Nelson LD, Simonsohn U. False-positive psychology: undisclosed flexibility in data collection and analysis allows presenting anything as significant. Psychol Sci. 2011;22(11):1359–66. 10.1177/0956797611417632

69. Vieth B, Parekh S, Ziegenhain C, Enard W, Hellmann I. A systematic evaluation of single cell RNA-seq analysis pipelines. Nat Commun. 2019;10(1):4667. 10.1038/s41467-019-12266-7

70. Fang C, Selega A, Campbell KR. Beyond benchmarking and towards predictive models of dataset-specific single-cell RNA-seq pipeline performance. Genome Biol. 2024;25(1):159. 10.1186/s13059-024-03304-9

71. Hie B, Bryson B, Berger B. Efficient integration of heterogeneous single-cell transcriptomes using Scanorama. Nat Biotechnol. 2019;37(6):685–91. 10.1038/s41587-019-0113-3

72. Wegmann R, Neri M, Schuierer S, Bilican B, Hartkopf H, Nigsch F, et al. CellSIUS provides sensitive and specific detection of rare cell populations from complex single-cell RNA-seq data. Genome Biol. 2019;20(1):142. 10.1186/s13059-019-1739-7

73. Liu S, Thennavan A, Garay JP, Marron JS, Perou CM. MultiK: an automated tool to determine optimal cluster numbers in single-cell RNA sequencing data. Genome Biol. 2021;22(1):232. 10.1186/s13059-021-02445-5

74. Nguyen HCT, Baik B, Yoon S, Park T, Nam D. Benchmarking integration of single-cell differential expression. Nat Commun. 2023;14(1):1570. 10.1038/s41467-023-37126-3

75. Holland CH, Tanevski J, Perales-Paton J, Gleixner J, Kumar MP, Mereu E, et al. Robustness and applicability of transcription factor and pathway analysis tools on single-cell RNA-seq data. Genome Biol. 2020;21(1):36. 10.1186/s13059-020-1949-z

76. Patterson-Cross RB, Levine AJ, Menon V. Selecting single cell clustering parameter values using subsampling-based robustness metrics. BMC Bioinformatics. 2021;22(1):39. 10.1186/s12859-021-03957-4

77. Heumos L, Schaar AC, Lance C, Litinetskaya A, Drost F, Zappia L, et al. Best practices for single-cell analysis across modalities. Nat Rev Genet. 2023;24(8):550–72. 10.1038/s41576-023-00586-w

